# De novo Protein Generating Model via Structure-Aware Graph Neural Networks

**DOI:** 10.64898/2026.10.27.762190

**Authors:** Chance Gammill, Lakshman Tamil

## Abstract

The focus of this paper is to create a protein conformation prediction model that is optimized for structural accuracy rather than sequence recovery. To achieve this goal, we created a graph neural network (GNN) [19] for the design of computational proteins that integrates differentiable structure prediction through ESMFold [13] via a structural loss function, allowing direct optimization of the 3D conformational accuracy. Our model takes protein backbone coordinates as input and predicts an amino acid sequence using a composite loss that heavily weights structural similarity. On a matched evaluation sample, our model achieves structural precision (RMSD, TM-score, and GDT-TS) that is statistically indistinguishable from ProteinMPNN, a leading sequence recovery-focused method, despite training with no direct sequence recovery supervision and substantially lower native sequence recovery. Our results demonstrate that sequence recovery is not necessary to achieve competitive structural accuracy and that proteins can possess multiple non-unique solutions, suggesting that protein sequence prediction models may perform comparably well under purely structural conformation training. This distinction in training data and model structure allows for the training of a protein-generating model on non-natural proteins. Creating a model designed for de novo protein generation.

## 1. Introduction

The use of artificial intelligence in protein structure prediction was popularized by AlphaFold [10] in 2020, in the Critical Assessment of protein structure prediction [17], demonstrating that deep learning models could predict a protein’s three-dimensional conformation from its amino acid sequence with near experimentally comparable results. This success raised the question: Given a target backbone structure, is it possible to predict an amino acid sequence that will fold into that conformation? Models such as ProteinMPNN [5] have made great strides in this inverse protein design problem, achieving up to a 52.4% sequence recovery rate. However, sequence recovery may not be the only determining factor in protein structure folding. This motivated the use of a model that directly optimizes for structural accuracy rather than sequence recovery. As de novo protein generation attempts to create protein sequences that may or may not exist in nature, using known amino acid sequences to train the model directly conflicts with the intended goal. So, this paper attempts to create a model that trains without using an amino acid sequence.

## 2. Data Accessibility

All Protein structures in this work were obtained from the Protein Data Bank (PDB) [3] using an AlphaFold3 [1] `fetch_databases.sh` function, which retrieves the experimentally determined structures in mmCIF format. Each structure was decomposed into individual chains and preprocessed into backbone coordinate representations consisting of N, C*α*, C, and O atomic positions. All available chains were used regardless of native length and missing residues, then truncated to a fixed length of 128 residues. Any chain shorter than 128 residues was padded with a placeholder “unknown residue” token. This truncation strategy was judged to reduce, but not eliminate, redundancy between the training and test partitions, since fixed-length fragments drawn from different regions of different proteins are structurally and compositionally more diverse than their full-length source sequences even when the source proteins are homologous. For this reason, no additional sequence identity or structural similarity clustering was applied. It should also be noted that this work trains the sequence design model directly against ESMFold-predicted structures; therefore, any of ESMFold’s own pretraining biases will inevitably show up in the GNN model. As ESMFold’s own weights are kept frozen throughout training and evaluation, any biases in the ESMFold model cannot be completely controlled or eliminated and are acknowledged as a limitation of relying on a pretrained structure predictor for both training supervision and evaluation. Additionally, the underperformance of the normally high-performing ProteinMPNN model on this data set is expected, as a result of the truncated sequences; this is examined in Section 4.2. This data set was chosen specifically to allow the models and testing data to exceed the known protein folding structures on which the compared models were originally trained and validated.

The data pull was performed in February 2026 and retrieved 200,000 PDB structures for the training data set and another 10,000 structures for evaluation. No filtering criteria beyond the fixed 128-residue truncation and padding described above were applied to either dataset; structures were not excluded by resolution, experimental method, or chain type. No special handling was applied for missing backbone atoms in the source coordinates, and any gaps present in the deposited structures were passed through as is. Of the 10,000 evaluation structures pulled, a matched comparison was completed between all evaluated methods for 224 proteins, a practical limit given the computational cost of running the full evaluation pipeline per target.

No separate held-out validation partition was carved out from the training pool; the 200,000-structure training set and 10,000-structure test set described above are the only two partitions used. Because training was conducted over ten sessions drawn from different subsets of training data, the model did not see an individual protein more than twice during the full training process, limiting the risk of overfitting to any single structure that a typical validation-based stopping criterion would otherwise prevent.

## 3. Methods

We utilize a structure-aware graph neural network (GNN) [19] for computational protein design, which takes a protein backbone structure as input and predicts optimized amino acid sequences for structural replication, optimized through a translational loss function. Instead of comparing our predicted sequence with the true sequence, we predict the structure into which the sequence would fold, using ESMFold [13], and compare it with the original structure, enabling backpropagation without exposing the model to any sequences. To make this possible, our architecture follows an encoder-decoder framework inspired by ProteinMPNN [5], augmented with a differentiable structure prediction module utilizing ESMFold [13], which enables structural supervision during training.

### 3.1. Model Architecture and Training

The key distinction between our approach and existing methods such as ProteinMPNN is the training objective and process. ProteinMPNN [5] focuses on achieving a high sequence recovery rate by optimizing the cross-entropy between predicted and native amino acid sequences. Although this produces high sequence recovery rates, it does not necessarily mean that the designed sequence will fold into the desired protein structure; this approach also has limitations when it comes to the generation of new proteins, as these types of models can only be trained on existing proteins and therefore lack the fundamental ability to create new sequences. Our model addresses these limitations by incorporating a differentiable structural loss function, where predicted amino acid probabilities are supplied to ESMFold via Gumbel-Softmax reparameterization [8], and the resulting predicted coordinates are compared against the ground truth backbone. This allows for optimization of the structural quality of the predictions, rather than the sequence recovery. This also means that any desired shape could be intentionally overtrained if someone so pleased, as the only necessary input is the structure.

#### 3.1.1 Graph Representation

Each protein sequence backbone structure is modeled as a *k*-nearest neighbor graph (k-NN) [4], 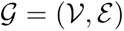, where the nodes 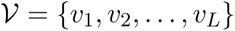 correspond to the residues, and the *E* edges connect each residue to its *K* nearest spatial neighbors based on the distance of C*α* in the 3-D space. Each residue is represented by the Cartesian coordinates of four backbone atoms N, C*α*, C, and O, which generate an input tensor **X** *∈* ℝ*^L^*^×*A*×*C*^. In our model, *K* = 20, *L* is the length of the sequence, *A* is the number of backbone atoms, and *C* is the number of Cartesian coordinates.

#### 3.1.2 Radial Bias Function

Prior to feature extraction, backbone coordinates are aligned with a three-point local reference frame: the C atom of the first residue is placed at the origin, the C atom of the second residue is placed on the positive *x*-axis, and the N atom of the first residue is placed in the *xy*-plane with positive *y*. This alignment is computed through sequential translation and Rodrigues’ rotation [18], providing a standard location and orientation, reducing sensitivity to arbitrary global positions. This pre-processing enables rotational invariance, simplifying downstream learning. Alternatively, equivariance can be built into the network itself through geometric vector perceptron layers that operate directly on vector-valued features [9].

#### 3.1.3 Edge Features

Edge features encode the sequential and spatial relationships between residues. The spatial proximity is captured using a radial basis function (RBF) encoding of pairwise distances, given in Equation 1:

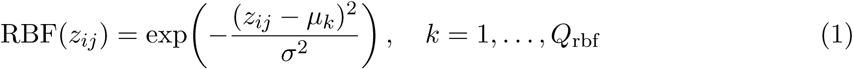

where 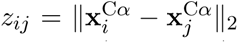 is the Euclidean distance between the C*α* atoms of residues *i* and *j*, and *µ_k_* are (*Q*_rbf_ = 16) centers linearly spaced between *D*_min_ = 2 Å and *D*_max_ = 22 Å, and *σ* = (*D*_max_ *− D*_min_)*/Q*_rbf_.

Sequential relationships are encoded using 16 relative positional embeddings that represent the offset of the clipped residue index Δ*ij* = *j − i* between the connected nodes of the intrachain edges. The concatenation of the RBF features from Equation 1 and the positional features is projected through a linear layer followed by layer normalization [2], producing the final edge embeddings 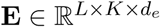, where *L* = length of sequence, limited to a maximum of 128, *K* = 20 is the number of neighbors per residue, and *d_e_* is the edge feature dimension.

Although not in this model, it is important to acknowledge that the model could possibly be improved by accounting for the other backbone atoms. Such a method would allow the model to improve its rotational understanding. However, such a change increases the final edge embeddings to 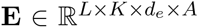. This drastically increases training and computational costs.

#### 3.1.4 Edge Featurization Pipeline

The complete pipeline is summarized in Figure 1. The coordinates of the residue backbone are first expressed in the local reference frame of Section 3.1.2. A *k*-nearest-neighbor graph over C*α* positions (*K* = 20) then defines the edge set, as described in Section 3.1.1. Each edge carries two complementary descriptors: a radial basis function encoding of the pairwise C*α* distance *z_ij_* (Equation 1), shown schematically in the inset of Figure 1, and a relative positional embedding of the clipped residue index offset Δ*_ij_*. Here, the spatial index offset Δ*_ij_* represents spatial proximity, and the RBF represents sequential adjacency, so the two descriptors jointly express where a neighboring residue sits in space and along the chain. The two are concatenated and projected through a linear layer with layer normalization to give the edge embeddings 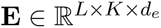, as mentioned in 3.1.3.

**Figure 1:**
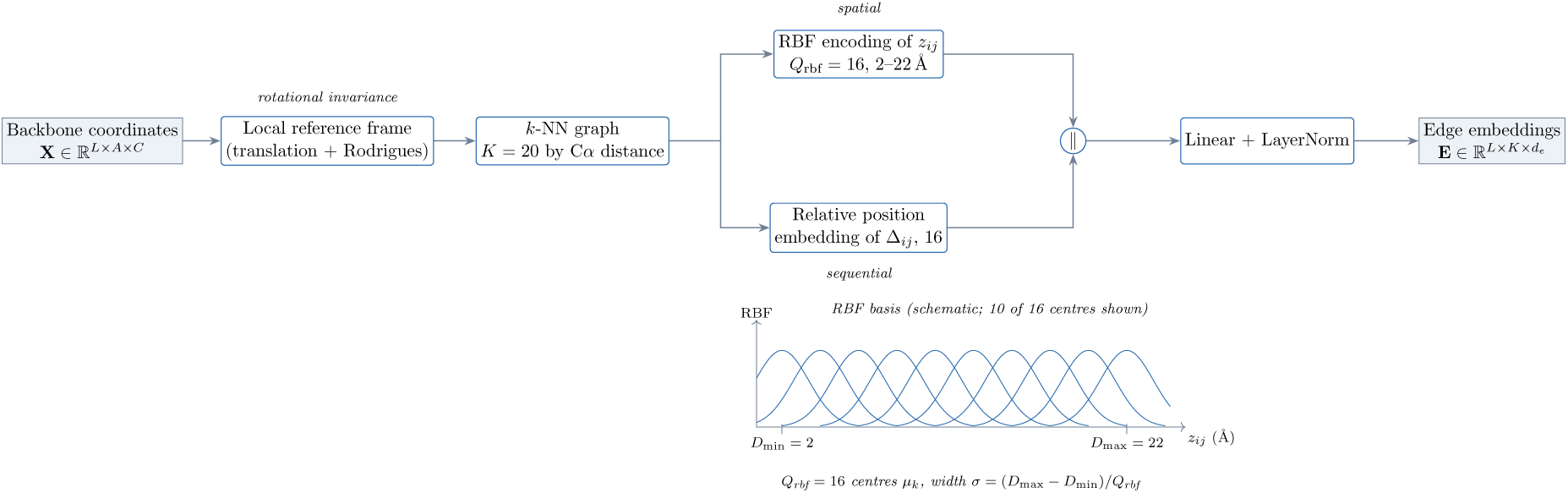
Construction of the edge features, from backbone coordinates to the edge embeddings 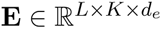, with the radial basis shown schematically in the inset.

#### 3.1.5 Encoder

The encoder consists of *N*_enc_ = 4 bidirectional message passing layers [6] which update the node and edge representations; the structure of a single such layer is shown on the left of Figure 2. In each layer *l*, the node features are updated by aggregating features from the neighboring edges and nodes, as given in Equation 2:

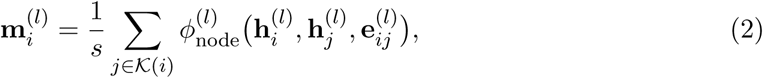

**Figure 2:**
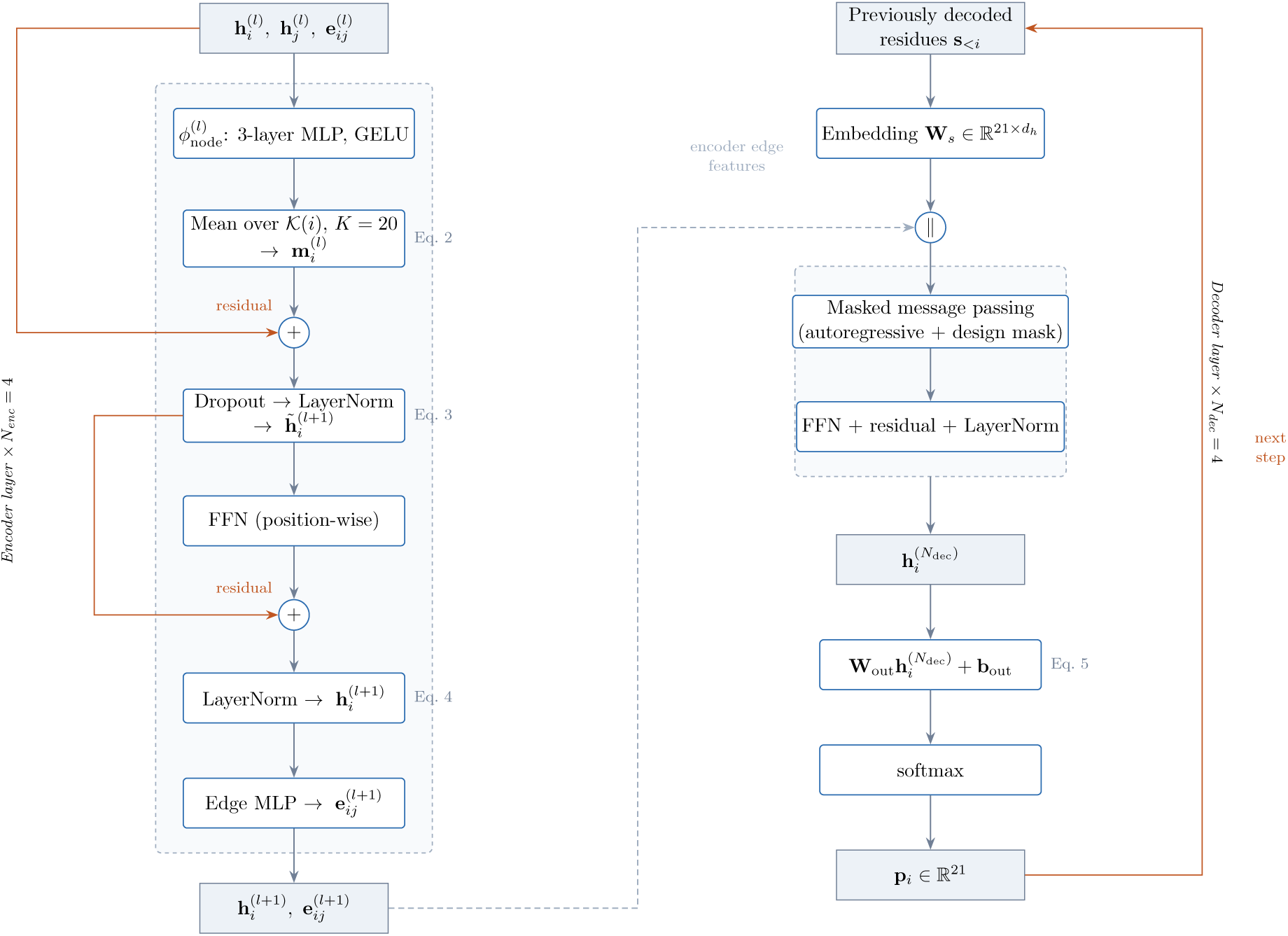
Structure of a single encoder layer (left) and the decoder (right), with orange paths marking the residual and autoregressive feedback connections.

where 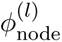 is a three-layer multilayer perceptron (MLP) with GELU activation [7], 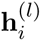 is the feature vector of node *i* at layer 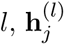 is the feature vector of the neighboring node, 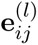 is the edge feature vector from node *i* to *j*, *s* is the normalization constant, *K*(*i*) denotes the *K*-nearest neighbors of node *i*, and 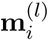 represents the aggregation of all of this information to update the node *i*’s representation.

The node features are then updated using a residual connection, followed by layer normalization, as given in Equation 3:

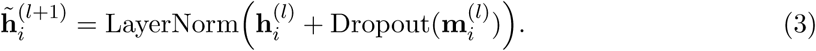

Each node update is followed by a position-wise feed-forward network (FFN) with a residual connection, as given in Equation 4:

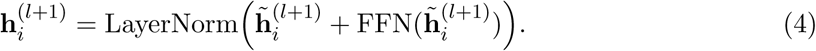

Edge features are then updated using an MLP that takes the updated features as input for the connected node pair, shown as the final stage of the encoder column in Figure 2. This bidirectional update of the edges and nodes allows backpropagation along the protein backbone and spatial contacts.

#### 3.1.6 Decoder

The decoder auto-regressively generates the amino acid sequence, conditioned on the structural encoding; its layout is shown on the right of Figure 2. In each decoding step, the identities of the previously predicted residues are embedded using a learned embedding matrix 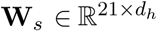, where *d_h_* is the hidden dimension, and 21 represents the 20 amino acids and the 1 unknown token. The resulting per-residue sequence embeddings are concatenated with the corresponding encoder-derived edge features. The autoregressive mask applies only to the amino acid identities: the sequence embedding is zeroed at every position not yet decoded, so that the non-decoded positions only contribute structural features. The encoder-derived node and edge features 3.1.5 are never masked and remain visible at every position regardless of the decoding order. Each residue is therefore conditioned on the complete backbone geometry, but only on the subset of amino acid identities decoded before it.

The decoder consists of *N*_dec_ = 4 layers, each performing masked message passing under the combined autoregressive and design mask indicated in Figure 2, followed by an FFN with residual connections and layer normalization as in the encoder of Section 3.1.5. The hidden representation produced by the final decoder layer is then projected and normalized into a per-residue distribution over the 21 tokens, as given in Equation 5:

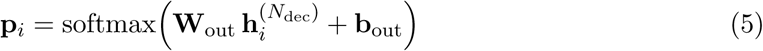

where 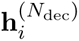 is the hidden representation of residue *i* after the final decoder layer, **W**_out_ is a learned output projection matrix that maps this hidden representation to a non-normalized score for each of the 21 residue tokens, and **b**_out_ is a learned bias vector that adds a per-token offset to those scores. This produces a probability distribution **p***_i_*, which is used to predict the amino acid at that residue in the autoregressive sequence.

#### 3.1.7 Encoder and Decoder Structure

Figure 2 summarizes a single encoder layer alongside the decoder. Each of the *N*_enc_ = 4 encoder layers aggregates messages from the *K* nearest neighbors of residue *i* through a three-layer MLP with GELU activation (Equation 2), applies a residual connection with dropout [20] and layer normalization (Equation 3), and is followed by a position-wise feed-forward network under a second residual connection (Equation 4); a separate MLP then updates the edge representations. The decoder embeds previously decoded residue identities through **W***_s_*, concatenates them with the encoder-derived edge features, and applies *N*_dec_ = 4 layers of masked message passing under a combined autoregressive and design mask. The final hidden state is projected by **W**_out_ and normalized by a softmax to give the per-residue distribution **p***_i_* (Equation 5). The next residue identity is then the highest probability in **p***_i_*.

In Figure 2, the orange paths mark the residual and autoregressive feedback connections, and the dashed path marks the encoder-derived edge features passing into the decoder; this transfer keeps the full backbone geometry visible at every decoding position regardless of the order in which residues are decoded.

The depth of each stack determines how much structural context informs a residue. A single encoder layer updates residue *i* using only the *K* = 20 neighbors in *K*(*i*), representing its immediate spatial environment. Stacking layers widens this receptive field; the second layer **h***_i_* has received information from the neighbors of its neighbors, and after *N*_enc_ = 4 layers it draws on residues up to four edges away in the *k*-NN graph. Because the *k*-NN graph is built on spatial proximity rather than sequence adjacency, these neighbors traverse the folded structure rather than sequential neighbors, so a residue accumulates context from positions that are sequentially distant but tightly packed.

#### 3.1.8 Differentiable Structure Prediction

To incorporate structural feedback during training, we integrated ESMFold [13] as a differentiable structure prediction oracle model. ESMFold v1 (`facebook/esmfold_v1`) was incorporated through the Hugging Face `transformers` library (`EsmForProteinFolding, EsmTokenizer`), run with full float32 precision for numerical stability. ESMFold’s own weights were kept frozen throughout training, so only the GNN’s parameters were updated; ESMFold itself served purely as a fixed differentiable critic. Prior work has approached this supervision differently; AlphaFold distillation trains a lightweight surrogate to mirror the structural quality estimates, avoiding the cost of a folding passes during sequence design [16]. We instead retain the folding model itself inside the training loop, accepting the added cost in exchange for gradients that pass through the predicted structure rather than through an approximation of it. The key challenge is that ESMFold expects discrete token inputs, which are non-differentiable. We addressed this issue by using Gumbel-Softmax reparameterization [8], given in Equation 6:

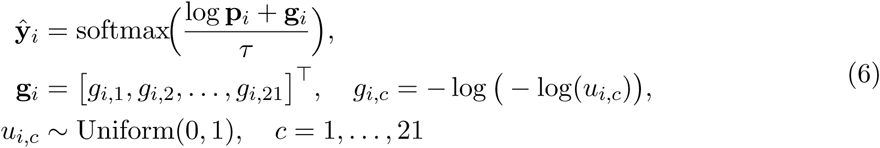

where *τ* = 1 is a temperature parameter controlling the sharpness of the approximation, *g_i,c_* is the Gumbel noise, *u_i,c_* is a random uniform variable, *i* is the residue, and *c* is the current token (Amino Acid). The resulting soft one-hot vectors **ŷ***_i_* are converted to continuous ESMFold embeddings via a weighted sum over the token embedding matrix, allowing gradients to flow from the predicted structure back through the sequence prediction network. This differentiable path, and the separate non-differentiable arg max branch used only for reporting, are shown in Figure 3. This enables the structural loss metric, given as 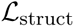 in Equation 7 and described in Section 3.1.9, to directly impact the model’s sequence predictions. The folded path is not the only route by which the predicted sequence is penalized. Of the five terms in the training objective, only the Gumbel cross-entropy term requires a folding pass; the structural proxy, repetition, diversity, and contact terms are all computed directly from the predicted amino acid probabilities, scoring the distribution itself against backbone geometry derived from the ground-truth coordinates. The predicted sequence therefore receives gradient signal at every step from several independent directions, of which the folded structure is the most expensive and the most directly tied to the design objective.

**Figure 3:**
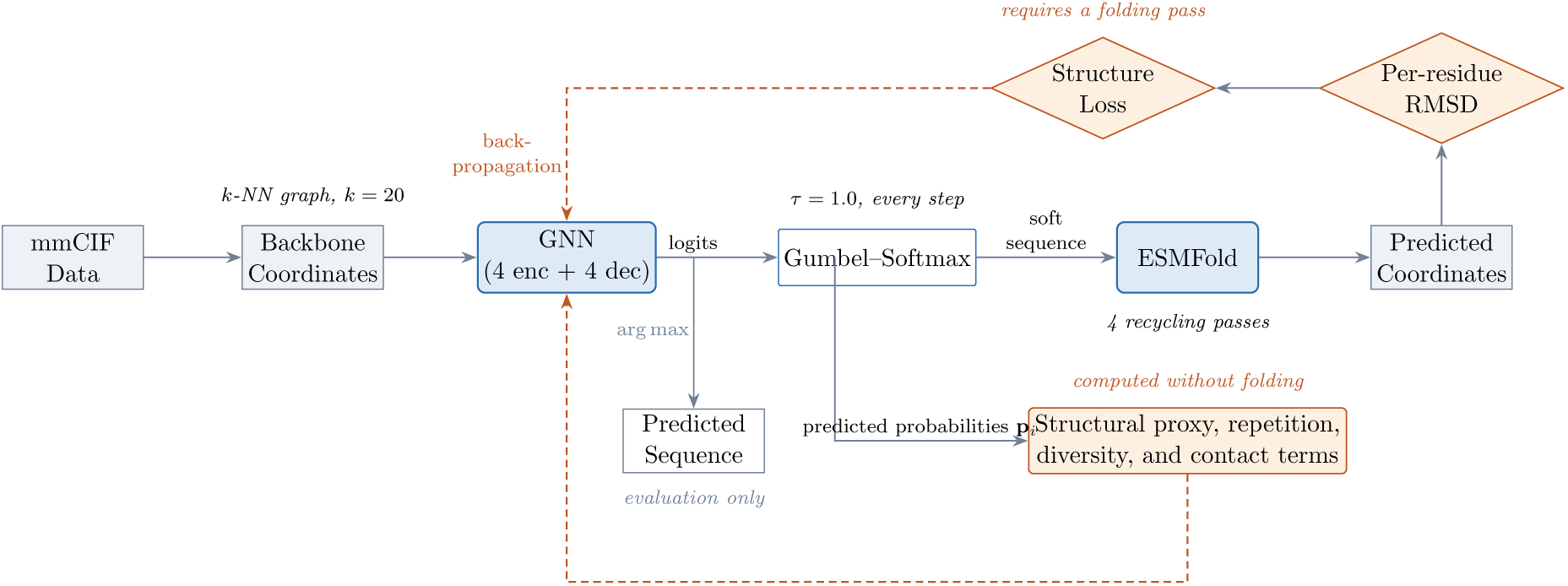
Overview of the structure-aware graph neural network and its differentiable training loop, showing both the folded path that supplies the structural signal and the loss terms computed directly from the predicted probabilities.

ESMFold was run with 4 recycling passes throughout training. The “unknown residue” padding token was excluded from the sequence passed into ESMFold at the embedding boundary rather than being assigned a soft embedding; padded positions were therefore never folded by ESMFold, though they remain represented as placeholder positions in downstream per-residue error tensors aligned to the fixed 128-residue representation. No formal validation-based checkpoint selection criterion was used; the checkpoint used for all reported results is the furthest trained checkpoint from training, since the goal was to expose the model to every sequence in the training set at least once rather than to optimize against a held-out validation metric. No random seed was fixed for training. Training was run on a single NVIDIA RTX 4080 Super GPU.

#### 3.1.9 Training Objective

Our model is trained using a composite loss function combining a fast, per-step structural proxy with the slower but more accurate prediction obtained by differentially folding the model’s own predicted sequence with ESMFold at every training step, as given in Equation 7:

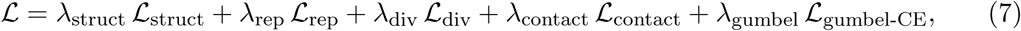

The five terms of Equation 7 are described below.

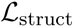 is a fast, per-step structural proxy computed directly from the model’s predicted amino acid probabilities. Without folding, it scores the expected helix/sheet propensity and hydrophobicity implied by the predicted sequence and simple backbone geometry features (local curvature and residue compactness) computed from the ground-truth coordinates; encouraging regions to favor helix/sheet-forming residues, straight regions to disfavor them, and encased positions to favor hydrophobicity accordingly. This fast structural proxy helped the model when ESMFold was not confident and increased consistency of the training, vastly reducing early stages of training time.

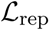 and 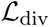 are a repetition penalty and a diversity penalty, respectively, which discourage the model from collapsing into repetitive or low-diversity amino acid sequences. This becomes especially important when increasing diversity, as using the same 4 amino acids is easier for the model early on.

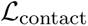 is an encasement loss function that encourages the predicted sequence to place hydrophobic residues at buried positions, independent of normal structure loss. A binary contact map is computed from the ground truth backbone, with *C_ij_* = 1 where the C*α* atoms of the residues *i* and *j* lie within 8.0 Å of each other and are otherwise zero. The expected hydrophobicity at each position is obtained from the model’s predicted amino acid distribution, meaning that it is trained from its own predictions and not from the original coordinates.

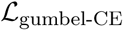 is the structural signal per residue, where at every training step, the probability of the predicted amino acids of the model is converted to a soft one-hot sequence via Gumbel-Softmax reparameterization (Equation 6) and differentially folded by ESMFold. The resulting per-residue RMSD between the predicted and ground-truth C*α* coordinates, which is used to reweigh a second cross-entropy term via hard-residue mining, so that residues with higher predicted structural error receive proportionally higher weights in the cross-entropy term. This allows the model to be penalized more for sequence choices at positions where its own predicted structure differed the most from the target backbone. This is different from sequence loss, as it is true structural error correction, concentrating gradient signals on structurally consequential positions rather than sequential differences, without needing information about the “True” sequence.

#### 3.1.10 Optimization

The model is optimized with Adam [12] with parameters *β*_1_ = 0.9, *β*_2_ = 0.98, *ɛ* = 10*^−^*^9^, and an initial learning rate of *η* = 10*^−^*^4^. A linear warm-up schedule was applied for the first 1,000 steps, followed by cosine annealing with periodic restarts (*T*_0_ = 1000, *T_mult_* = 2, *η_min_* = 0.1*×* the initial rate). Gradients were normalized to 1 for training stabilization. Gradients were accumulated over 4 mini-batches to increase the effective batch size. During training, Gaussian noise with standard deviation *ɛ_aug_* = 0.005 Å was added to the backbone coordinates to decrease susceptibility to volatile protein crystallography. Training was carried out over approximately 10 separate sessions of roughly 40,000 steps each, run over multiple different data sets/chunks, for a total of approximately 400,000 optimization steps.

#### 3.1.11 Model Summary

Figure 3 gives an overview of the structure-aware graph neural network and its differentiable training loop. Backbone coordinates extracted from mmCIF files are encoded as a *k*-nearest-neighbor graph over residues and decoded into per-residue logits. On the differentiable path, drawn solid, the logits are relaxed by a Gumbel-Softmax reparameterization into a soft sequence representation that ESMFold folds into predicted coordinates; per-residue RMSD against the native structure forms the structure loss, whose gradient, drawn dashed, propagates back through ESMFold to the network weights. The discrete sequence obtained by arg max, drawn in gray, is used only for reporting and evaluation and lies outside the gradient path. This separation is what allows the model to be scored on the structure its sequence produces while never receiving a gradient through a discrete sequence. For clarity the figure shows only the structural term of the training objective; the full objective is given in Equation 7.

Table 1 summarizes the key architectural and training hyper-parameters.

**Table 1:**
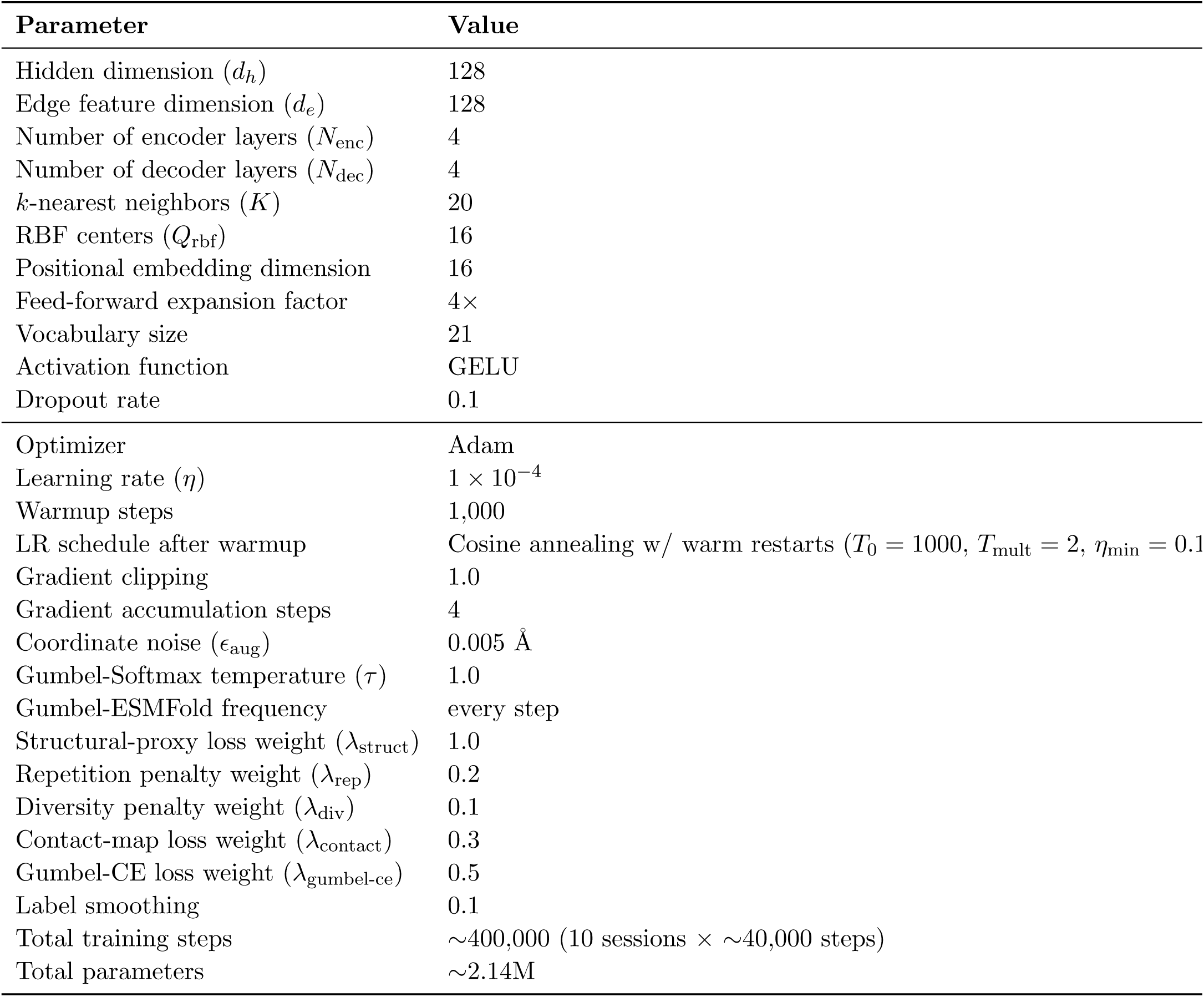
Model architecture and training hyper-parameters.

| Parameter | Value |
| --- | --- |
| Hidden dimension ($d_h$) | 128 |
| Edge feature dimension ($d_e$) | 128 |
| Number of encoder layers ($N_{\text{enc}}$) | 4 |
| Number of decoder layers ($N_{\text{dec}}$) | 4 |
| $k$-nearest neighbors ($K$) | 20 |
| RBF centers ($Q_{\text{rbf}}$) | 16 |
| Positional embedding dimension | 16 |
| Feed-forward expansion factor | $4\times$ |
| Vocabulary size | 21 |
| Activation function | GELU |
| Dropout rate | 0.1 |
| Optimizer | Adam |
| Learning rate ($\eta$) | $1 \times 10^{-4}$ |
| Warmup steps | 1,000 |
| LR schedule after warmup | Cosine annealing w/ warm restarts ($T_0 = 1000$, $T_{\text{mult}} = 2$, $\eta_{\text{min}} = 0.1$) |
| Gradient clipping | 1.0 |
| Gradient accumulation steps | 4 |
| Coordinate noise ($\epsilon_{\text{aug}}$) | 0.005 Å |
| Gumbel-Softmax temperature ($\tau$) | 1.0 |
| Gumbel-ESMFold frequency | every step |
| Structural-proxy loss weight ($\lambda_{\text{struct}}$) | 1.0 |
| Repetition penalty weight ($\lambda_{\text{rep}}$) | 0.2 |
| Diversity penalty weight ($\lambda_{\text{div}}$) | 0.1 |
| Contact-map loss weight ($\lambda_{\text{contact}}$) | 0.3 |
| Gumbel-CE loss weight ($\lambda_{\text{gumbel-ce}}$) | 0.5 |
| Label smoothing | 0.1 |
| Total training steps | $\sim 400,000$ (10 sessions × $\sim 40,000$ steps) |
| Total parameters | $\sim 2.14\text{M}$ |

### 3.2. Evaluation Metrics

We evaluated the structural accuracy of the predicted and native sequences using a wide range of metrics, mainly by folding the sequences using ESMFold [13] to obtain a three-dimensional coordinate predicted structure, which was then compared with the ground-truth backbone structures after Kabsch alignment [11].

#### 3.2.1 Global Structural Metrics

##### Root-Mean-Square Deviation (RMSD)

We computed RMSD [15] between C*α*s using the ESMFold-predicted structure of the predicted sequence and the ground-truth backbone after Kabsch superposition, as given in Equation 8:

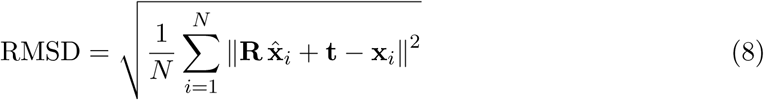

where **x̂***_i_* and **x***_i_* denote the predicted and native C*α* coordinates of residue *i*, respectively, **R** and **t** are the optimal rotation and translation obtained via Kabsch alignment, and *N* is the number of residues included in the comparison.

##### TM-score

The template modeling score (TM-score) [23] is a length-normalized measure of structural similarity that is less sensitive to local outliers than RMSD, as given in Equation 9:

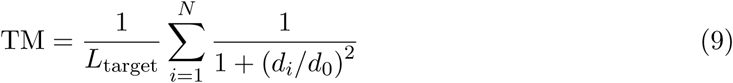

where *d_i_* is the distance between the *i*-th pair of aligned C*α* atoms after superposition, *N* is the number of aligned residues, and 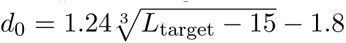 is a length dependent normalization factor. Statistical analysis of PDB structures indicates that scores below approximately 0.17 are characteristic of randomly chosen unrelated protein pairs, whereas scores above 0.5 generally correspond to pairs sharing the same fold in SCOP/CATH [21].

##### GDT-TS

The Global Distance Test–Total Score (GDT-TS) [22] measures the fraction of residues whose C*α* atoms fall within multiple distance thresholds after Kabsch alignment, as given in Equation 10:

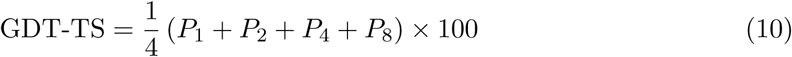

where *P_d_*denotes the percentage of residues whose C*α* atoms are within *d* Å of their native positions.

##### pLDDT

The predicted Local Distance Difference Test (pLDDT) [14], reported by ESMFold, provides a per-residue estimate of model confidence in the predicted local backbone geometry. Higher pLDDT values indicate greater confidence in the accuracy of local structural features, while lower values typically correspond to flexible or poorly resolved regions.

## 4. Results

Using the metrics defined in Section 3.2, we evaluated our structure-aware GNN model on a separate set of *n* = 224 evaluation proteins, not included in the training or testing data, using ESMFold to predict the structure of sequences from three categories: the native sequence, our model’s predicted sequence, and ProteinMPNN’s predicted sequence. We compared the resulting structures with the provided crystal structure coordinates after Kabsch superposition. Both our model and ProteinMPNN were evaluated with a matched best-of-5 sampling; the sequence with the lowest self-reported RMSD for both our model and ProteinMPNN was carried forward for structural comparison. Of the 224 evaluated proteins, 221 had valid results and were used for the paired statistical comparisons reported below.

### 4.1. Evaluation Metrics

The three primary structural metrics reported in this section are defined in Equations 8 and 10 of Section 3.2.1.

In the matched evaluation sample, where *n* = 221, our model achieved a mean RMSD of 8.097 Å (Figure 4), compared to 8.500 Å for ProteinMPNN and 6.280 Å for the native sequence. Our model also achieved a mean TM-score of 0.491 (versus 0.495 for ProteinMPNN and 0.606 for native) and a mean GDT-TS of 41.73 (versus 42.65 for ProteinMPNN and 54.19 for native). Paired *t*-tests (confirmed by Wilcoxon signed-rank tests) show no statistically significant differences between our model and ProteinMPNN on any of these three primary structural metrics (RMSD: *p* = 0.221; TM-score: *p* = 0.805; GDT-TS: *p* = 0.516), indicating that our structurally-optimized model achieves similar results when compared with Protein-MPNN, despite the two methods being trained toward fundamentally different objectives. The spread of each metric across the evaluation set is shown in Figure 5; the distributions for our model and for ProteinMPNN overlap substantially, so the agreement between the two methods holds across the evaluation set and is not an artifact of two dissimilar distributions sharing a mean.

**Figure 4:**
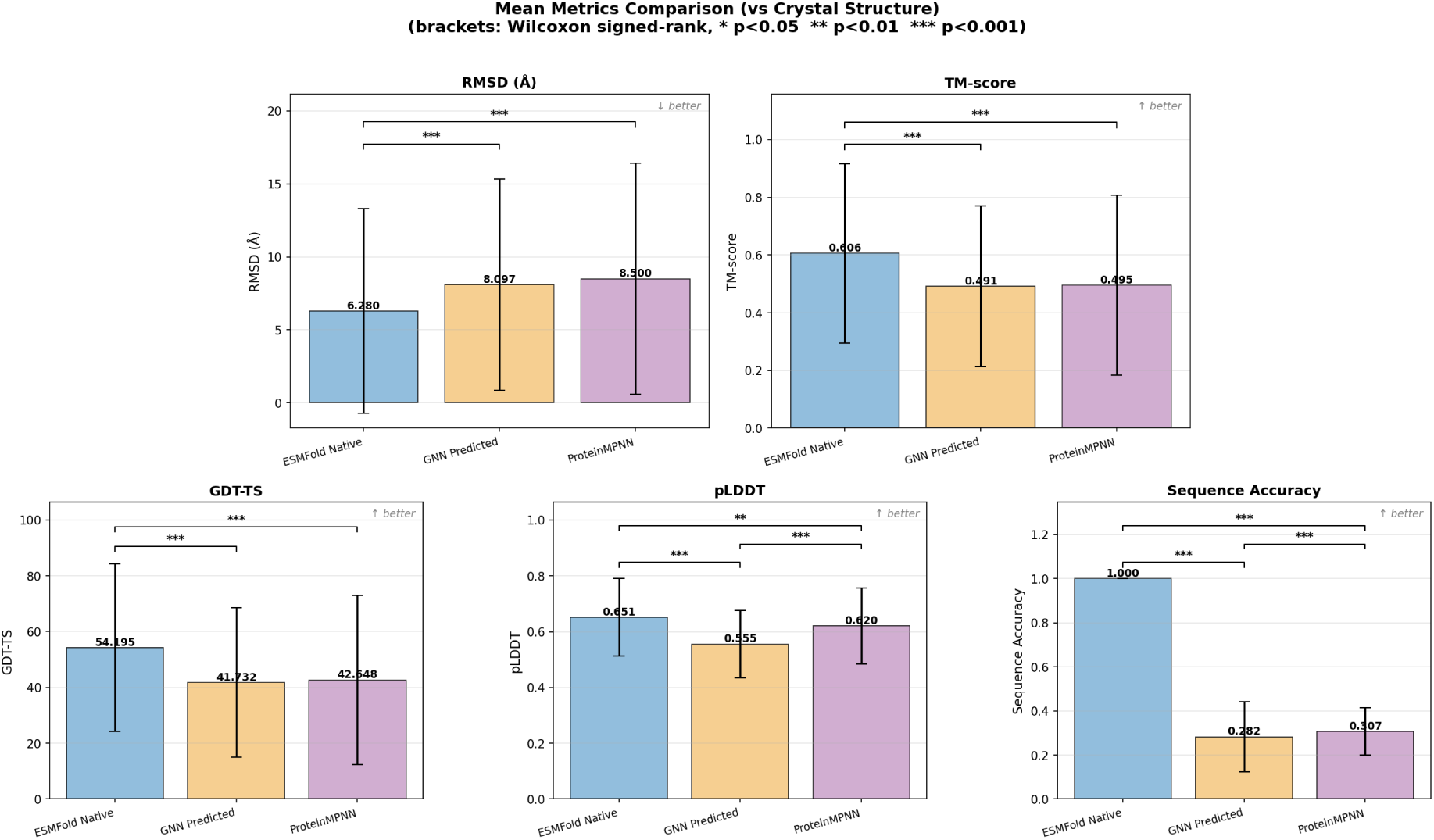
Mean structural comparison metrics across all evaluated proteins, Three sequence types are compared: the PDB native sequence, our model’s predicted sequence, and Protein-MPNN’s predicted sequence.

**Figure 5:**
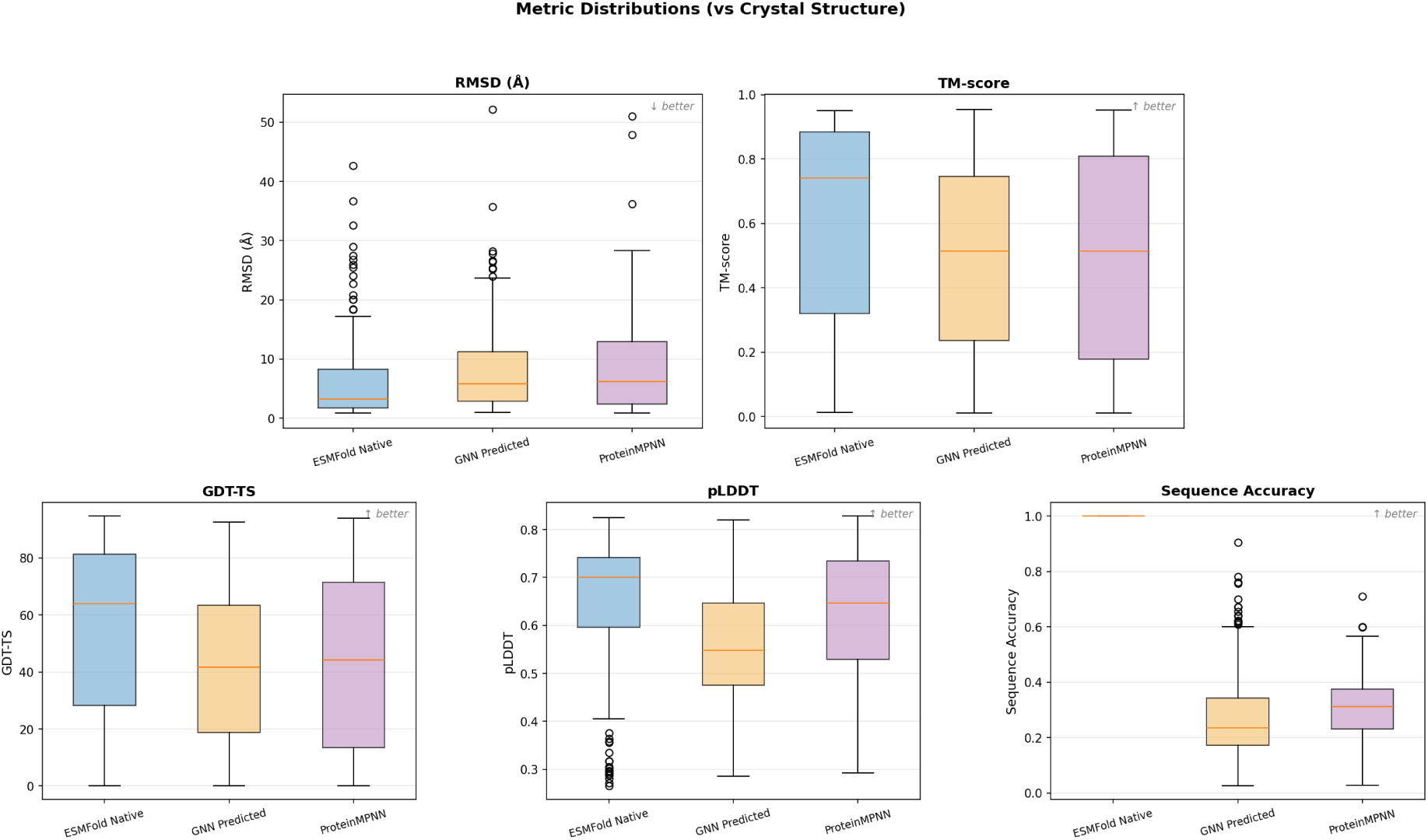
Distribution of structural comparison metrics across all evaluated proteins, shown as box plots giving the median, inter-quartile range, and spread for each metric. Three sequence types are compared: the PDB native sequence, our model’s predicted sequence, and ProteinMPNN’s predicted sequence.

This structural similarity comes with a lower sequence recovery, achieving a mean sequence accuracy of 0.282, significantly lower than ProteinMPNN’s 0.307 (*p* = 0.013). Both native and ProteinMPNN-designed sequences also achieved significantly higher mean pLDDT than our model (native: 0.651; ProteinMPNN: 0.620; ours: 0.555; both comparisons *p <* 0.001). These results are consistent with our model and are an expected outcome of producing de novo protein sequences, since generating sequences that deviate from the natural sequence space is expected to reduce ESMFold’s confidence. We do not treat the low pLDDT itself as evidence of successful novelty. However, this unpredictable nature is an inherent result of sequences that deviate from natural proteins.

### 4.2. Limitations and consequences of shortened chains

In (Figure 6), it can be seen that there are amino acid residues that, no matter how the sequence is changed, will always have a large error. This is likely a binding site for the protein’s tertiary structure, so it is possible that no shortened sequence would be capable of reproducing the provided protein structure.

**Figure 6:**
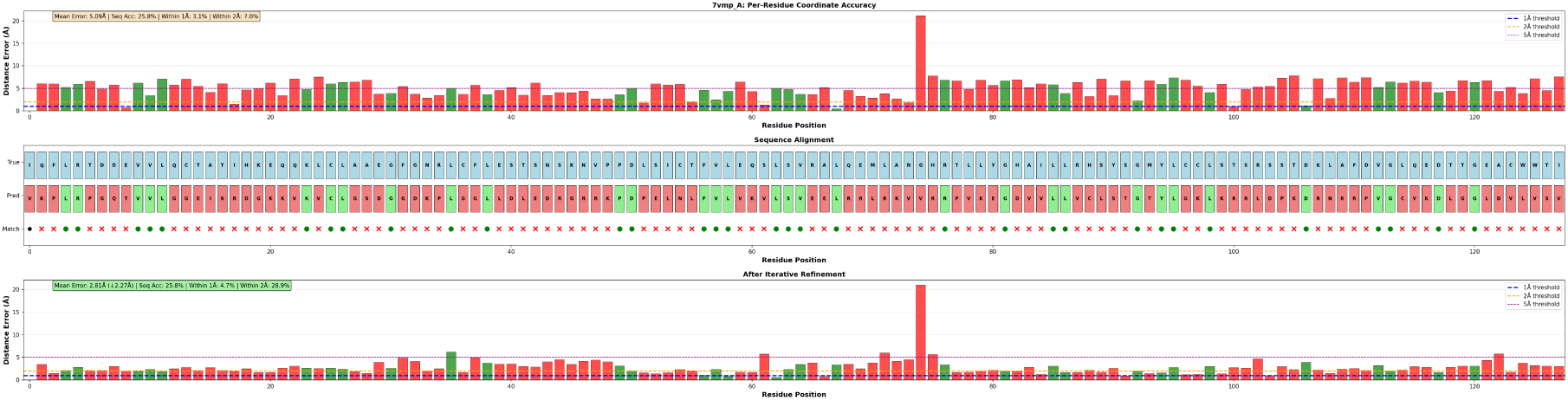
Per-residue C*α* distance error for the protein 7vmp. Displaying a low per residue distance error, but a single significantly flawed residue.

### 4.3. Non-unique solutions

Figure 7 shows three independent sampling attempts for the same target backbone, protein 7z4a. The three attempts (Figures 7a, 7b, and 7c) produce visibly different per-residue error profiles despite being drawn from the same model against the same structure, which demonstrates that the mapping from backbone to sequence is not one-to-one and that a single reported value conceals the range of outcomes the model can produce for one target. As reflected in these attempts, sequence recovery is not the only factor that affects the structural accuracy of a sequence. Additionally, Iteration C both our model and ProteinMPNN would likely achieve higher accuracies if the number of generations per sequence were increased further. This does not imply that any model would perform better with more attempts, but that the statistical results may underrepresent each model’s full potential. Both models make predictions and receive feedback within a short time frame using an already existing structure prediction model, which constraints allow for these models to make a large number of predictions with little cost.

**Figure 7:**
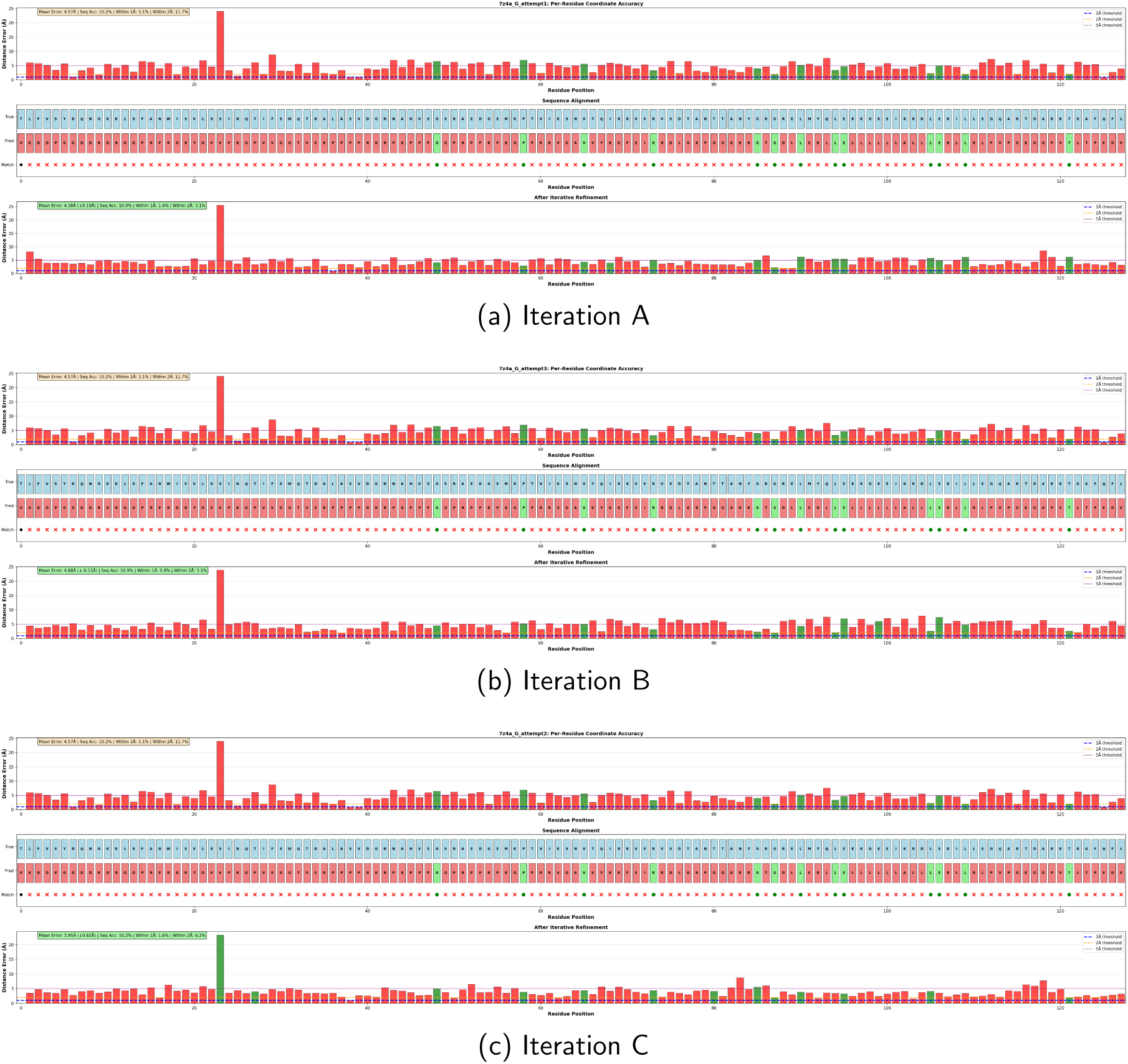
Per-residue C*α* distance error for three attempts of protein 7z4a, evaluating the same structure across different sampling attempts. Displaying the existence of non-unique solutions and how the mean does not fully reflect the range of outcomes a single model can produce.

### 4.4. Variance of Predictions

The per-residue error heatmaps (Figure 8) reveal that our GNN model and ProteinMPNN have different blind spots and some similar red zones. In many of the places where they possess blind spots, the same blind spots exist in the native sequence, but there are a few sections where it can be seen that our GNN model outperforms the native sequence. These heat maps provided ample evidence for a more in depth look into the models. This convinced us to run a per-protein graph, which showed where the models did better or worse than the native sequence.

**Figure 8:**
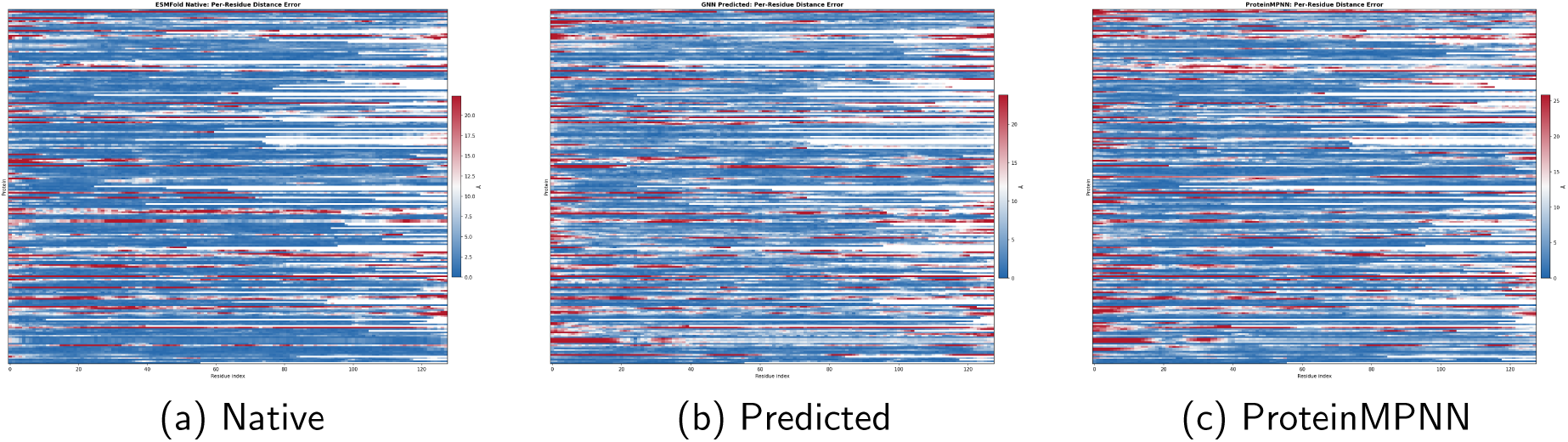
Per-residue structural error (Å) across all evaluated proteins (*n* = 224), aligned by sequence position, for each of the three sequence types.

### 4.5. Representative Per-Protein Case Studies

To illustrate the range of results summarized by the aggregate statistics in Figures 8 and Figure 4, we compiled per-residue error trajectories for three representative proteins: a case where our GNN model outperformed ProteinMPNN (7RS6, chain C), a case where our GNN model outperformed ProteinMPNN and the native sequence (7S0R, chain B), and a case where neither models were able to follow the native sequence (7S94, chain E).

These case studies illustrate a pattern similar to that observed in the heat maps of Section 4.4 (Figure 8). Our GNN model and ProteinMPNN do not always fail or succeed on the same proteins but instead possess different strengths and weaknesses. In 7RS6_C (Figure 9), our model’s predicted structure tracks the native backbone considerably more closely than ProteinMPNN’s across most of the sequence. Case 7S0R_B (Figure 10) is representative of a non-natural sequence that causes both the native sequence and the ProteinMPNN sequence to fail, but has no impact on the structurally focused model. Finally, 7S94_E (Figure 11) shows the reverse of the second case, with the ProteinMPNN’s prediction tracking the native structure more closely than our GNN model, but both models eventually failing.

**Figure 9:**
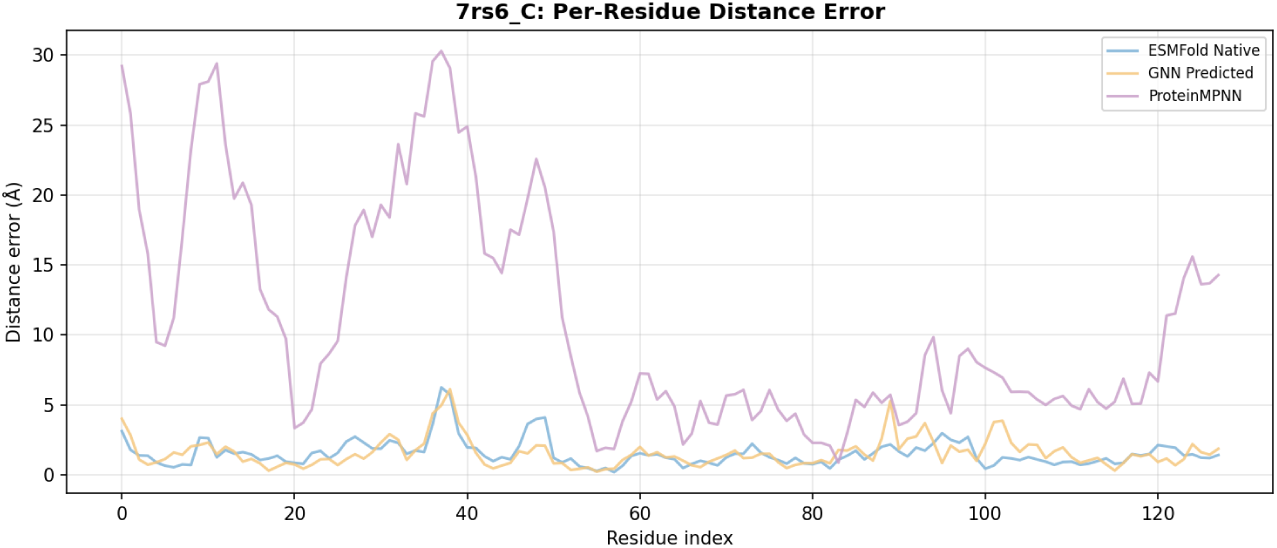
7RS6 C is a case where our GNN model closely followed the native sequence, while ProteinMPNN could not.

**Figure 10:**
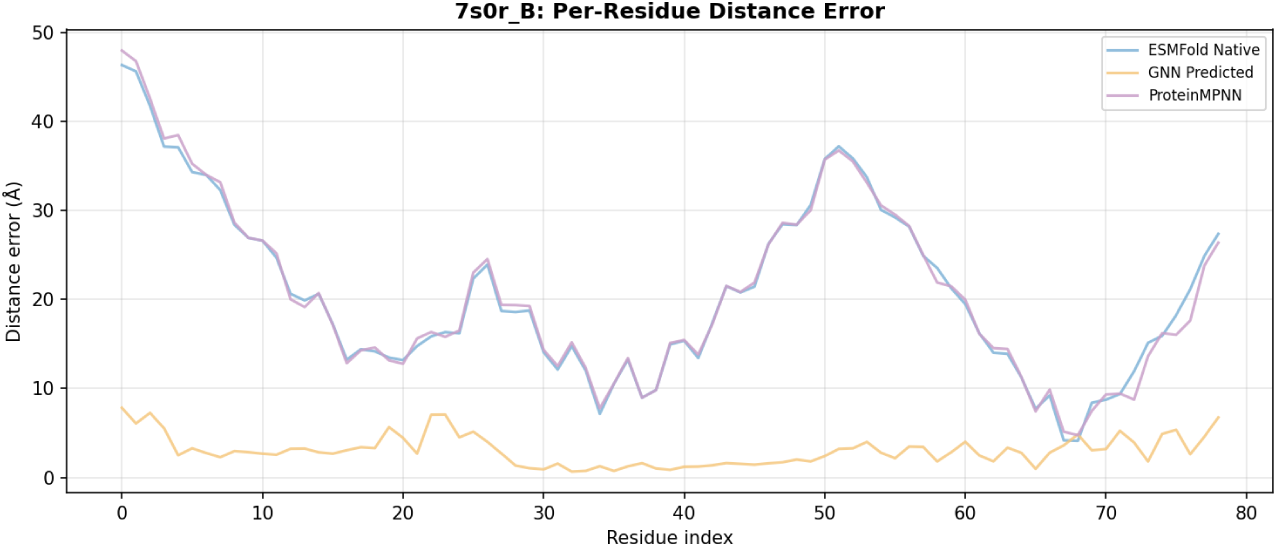
7S0R B is a case where our GNN model outperformed ProteinMPNN and the native sequence in per-residue structural error (Å).

**Figure 11:**
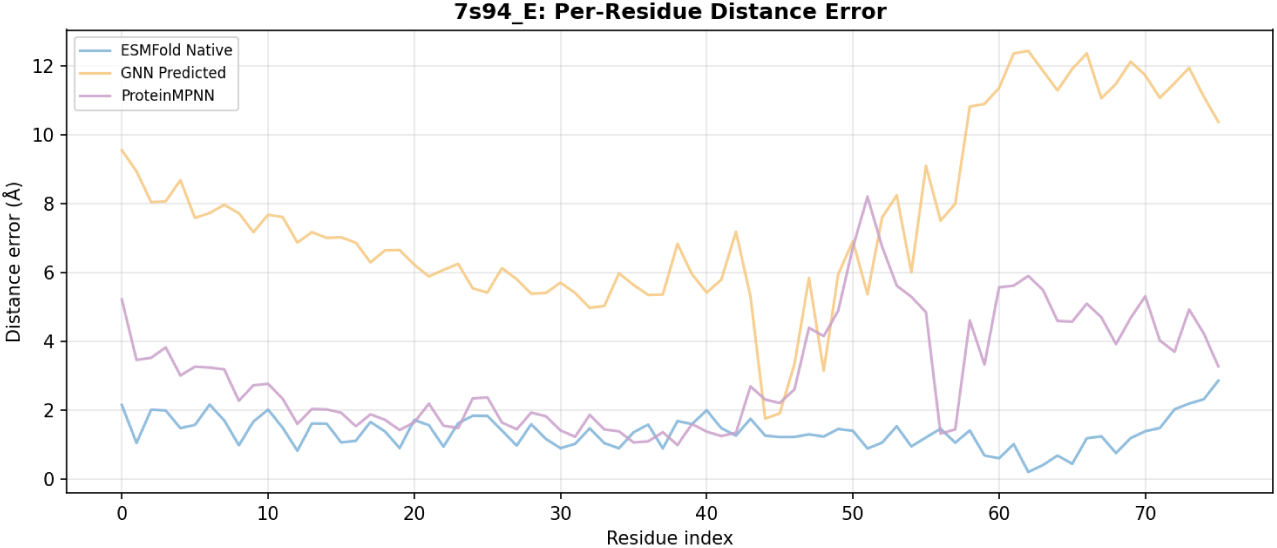
7S94 E displays a case where only the native sequence was able to stay accurate.

Taken together, these three cases indicate that the similarity observed in the aggregate statistics reflects two comparably capable methods that nonetheless make substantively different structural predictions on a per-protein basis.

## 5. Conclusion

In this paper, we presented a GNN for computational protein design that integrates differentiable structure prediction via ESMFold into the training loop, enabling end-to-end optimization with structure-level supervision in place of sequence-level supervision. We achieved this by back-propagating using Gumbel-Softmax and ESMFold, as described in Section 3.1.8 and given in Equation 6, allowing our model to directly optimize for structural replication rather than relying on sequence recovery. The composite objective that this enables is given in Equation 7 and described in Section 3.1.9, in which only the Gumbel cross-entropy term requires a folding pass.

Our results demonstrate that this structure-aware training approach achieves structural accuracy similar to ProteinMPNN, a leading sequence recovery-focused method, as measured by RMSD, TM-score, and GDT-TS on a matched evaluation sample of *n* = 221 proteins (Section 4.1). Our model reached a mean RMSD of 8.097 Å against 8.500 Å for Protein-MPNN, a mean TM-score of 0.491 against 0.495, and a mean GDT-TS of 41.73 against 42.65, with no statistically significant difference on any of the three metrics (RMSD: *p* = 0.221; TM-score: *p* = 0.805; GDT-TS: *p* = 0.516). This similarity was achieved despite our model receiving no native sequence supervision and possessing significantly lower sequence recovery than ProteinMPNN, at 0.282 against 0.307 (*p* = 0.013). All of this supports the conclusion that structural accuracy does not require close sequence replicability. We were also able to provide evidence that multiple sequences can be compatible with a single target backbone, shown across repeated sampling attempts on a single structure in Section 4.3, and that a model trained purely on structural similarity can match the structural performance of a sequence recovery-optimized model without needing to reproduce native sequences. The per-protein case studies of Section 4.5 further indicate that the two methods are not merely interchangeable: they succeed and fail on different targets, so the aggregate agreement reflects two comparably capable methods rather than two methods making the same predictions. Although not significantly different, our GNN model outperformed the ProteinMPNN model in mean RMSD by 0.403 Å, or 4.74%. Two limitations should be noted. Reliance on ESMFold as the structure prediction model during both training and evaluation means that errors or biases in ESMFold propagate to our model and to the comparison itself, and the fixed 128-residue truncation described in Section 2 both depresses ProteinMPNN’s performance relative to its published results and leaves some target backbones unreachable by any shortened sequence, as examined in Section 4.2.

In conclusion, we presented a structure-aware graph neural network for computational protein design that integrates differentiable structure prediction via ESMFold into the training process, enabling direct optimization of the 3-D conformational accuracy. This method allowed our model to match ProteinMPNN’s structural accuracy, a sequence-recovery-focused method, without negatively impacting the performance of the model. These results demonstrate that sequence recovery does not need to be the sole determining factor in model training and that there may be multiple unique solutions to a given native coordinate backbone. In addition, note that as protein structure prediction models, such as ESMFold, improve, so does the structure-aware GNN model.

## 6. Declarations

### Use of generative AI

Claude (Anthropic) was used for LaTeX formatting, editing, and drafting the TikZ source for Figures 1, 2, and 3. All figures were reviewed, corrected, and finalized by the authors. The authors assume full responsibility for the generated content. No generative AI tool was used to produce, analyze, or interpret the experimental results.

### Author contributions

C.G.: conceptualization, methodology, software, validation, formal analysis, data curation, writing – original draft. L.T.: supervision, writing – review & editing.

### Data availability

PDB accession identifiers and evaluation splits will be deposited with the archived code release.

### Code availability

Model code, evaluation scripts, trained weights, and other code are available at https://github.com/Justworkname/GenomeGenerator

### Ethics approval

Not applicable; this study used publicly available protein structures and did not involve human participants or animals.

